# Regime shifts in the Anthropocene: drivers, risks, and resilience

**DOI:** 10.1101/018549

**Authors:** Juan-Carlos Rocha, Garry D. Peterson, Reinette Biggs

## Abstract

Many ecosystems can experience regime shifts: surprising, large and persistent changes in the function and structure of ecosystems. Assessing whether continued global change will lead to further regime shifts, or has the potential to trigger cascading regime shifts has been a central question in global change policy. Addressing this issue has, however, been hampered by the focus of regime shift research on specific cases and types of regime shifts. To systematically assess the global risk of regime shifts we conducted a comparative analysis of 25 generic types of regime shifts across marine, terrestrial and polar systems; identifying their drivers, and impacts on ecosystem services. Our results show that the drivers of regime shifts are diverse and co-occur strongly, which suggests that continued global change can be expected to synchronously increase the risk of multiple regime shifts. Furthermore, many regime shift drivers are related to climate change and food production, whose links to the continued expansion of human activities makes them difficult to limit. Because many regime shifts can amplify the drivers of other regime shifts, continued global change can also be expected to increase the risk of cascading regime shifts. Nevertheless, the variety of scales at which regime shift drivers operate provides opportunities for reducing the risk of many types of regime shifts by addressing local or regional drivers, even in the absence of rapid reduction of global drivers.

## Introduction

We are living in the Anthropocene, an epoch where human actions intentionally and accidentally are changing planetary processes^1–5^ and ecosystems^6^. While some of these changes have been gradual, others have led to surprising, large and persistent ecological regime shifts^7,8^. Such shifts challenge ecological management and governance because they substantially alter the availability of ecosystems services^9^, while being difficult to predict and reverse^7^. While the importance of ecological regime shifts is increasingly recognized^3,10–12^, the variety of regime shifts and their drivers is less well known.

Following the exponential growth of the world’s economy, most drivers of global change are increasing^4,6,13^, and due to these increases the frequency and intensity of regime shifts are expected to increase too^14^. However most research on regime shifts is ill-suited to examine this proposition. Research on regime shifts has typically focused on theoretical models^8,15,16^, empirical evidence of regime shifts^17^, or potential early warnings signals^12,18^. These approaches require in-depth knowledge of the causal structure of the system or high-quality temporal data, leading to a focus on the analysis of particular cases of regime shifts. Here we complement this work by synthesizing and comparing different types of regime shifts in terms of global change impacts and opportunities for management. Our aim is to understand: What are the main drivers of regime shifts globally? What are their most common impacts on ecosystem services? And, what can be done to manage or avoid them?

## Materials and Methods

We addressed these questions using a diverse set of methods in a six phase process. First we developed a framework for data collection that facilitates comparison among regime shifts, namely the regime shifts database. Second, we identified and grouped the different drivers into hierarchical classes, distinguishing direct from indirect drivers. Third, strategies to manage regime shift drivers were identified and classified according to the scale at which action needs to be taken to tackle the effect of each driver. Fourth, to better understand what the main drivers of regime shifts are we studied their patterns of co-occurrence by constructing and simulating networks. Fifth, to discover what factors explained patterns among regime shifts and their drivers, exponential random graph models were used to explore what types of local interactions were consistent with the observed global patterns of the network. Sixth, to identify the most common impacts on ecosystem services, or the most common interactions among driver types, we analyzed the drivers and regime shifts datasets using ordering methods. Each of these steps are described in the following sections.

## Data

The regime shift database (RSDB) was created to synthesize, compare and share scientific knowledge about regime shifts in social-ecological systems [www.regimeshifts.org]. The RSDB currently provides a synthesis of >800 scientific papers, summarizing over 200 cases and about 25 generic types of regime shifts ^19^.It presents information both in plain text and 92 categorical variables about the i) main drivers of change, ii) impacts on ecosystem services, ecosystem processes and human well-being, iii) land use, ecosystem type and spatial-temporal scale at which each regime shift typically occurs, iv) possible managerial options, and v) assessment of the reversibility of the regime shift and the level of uncertainty related to the existence of the regime shift, and its underlying mechanism. The review of each regime shift is available online and wherever possible each entry has been written or peer-reviewed by an expert on the topic.

The database collects the most studied types of regime shifts in social-ecological systems^10^. Examples of regime shifts include i) well-established cases like eutrophication^17^, where lakes turn from clear water to murky water leading to reduced fishing productivity and toxic algae blooms; ii) controversial cases like dryland degradation when dry forest and savanna shift to deserts and bare soils, significantly reducing ecosystem services such as agricultural production and water cycling^20^; and iii) proposed shifts like the collapse of the Greenland ice sheet where the frequency and intensity of warm events will shift the ice sheet from permanent to occasional, reducing services such as coast line protection and climate regulation^21^. An overview of the 25 regime shifts analysed in this paper is given in S1 Table.

## Driver identification

Drivers include natural or human induced changes that have been identified as directly or indirectly producing a regime shift^6,22^. We first collected a preliminary list of drivers for each regime shift taking as a starting point that it should be referenced in the academic literature that the variable has causal influence on the regime shift.

For each regime shift we draw a causal loop diagram, a graphical representation of the causal structure of the system^23^. References and descriptions of each driver plus causal diagrams are available in the RSDB. To avoid ambiguities and conflicting definitions across different scholars, we defined drivers as variables outside the feedback mechanisms of the system, thus they are variables independent of the dynamics of the system. Direct drivers are those that influence the internal processes or feedbacks underlying a regime shift, and indirect drivers those that alter one or more direct drivers^22^. Based on the minimum distance to a feedback loop, we assessed the directedness of a driver as the shortest number of steps of separation to the feedbacks. This classification was done for each regime shift, therefore when comparing regime shifts a driver in one system can be part of an feedback in another.

To enable consisten comparison of drivers we systematically ensured that drivers were defined consistently across the database. After the first identification of drivers we checked for semantic cohesion, to avoid different words referring to the same driver. So for example cropping and agriculture were renamed agriculture. When the variables explicitly referred to different phenomena, different names were kept. For example rainfall variability and precipitation were kept separately as the first refers to variability and the second to total quantity. We further classified drivers as belonging to different types of global change by slightly modifying previous classifications^10,22^. We identified 15 detailed categories of drivers, which were further grouped into 5 broad categories: *habitat modification, food production, nutrients and pollutants, resource extraction* and *spill-over effects.* Thus, we distinguish between drivers stemming directly from human activities (e.g. fertilizer use) and drivers affected by the knock-on or ‘spill-over’ effects of these activities on natural processes (e.g. sedimentation or upwelling). A worked example is presented in S1 File.

## Scale of management

To examine management options for drivers of regime shifts we classified each driver by the scale it could be managed. Managerial options for each regime shift are synthesized in the RSDB. We exclusively classified each driver as requiring management at either local, national, or international scales. We considered a driver to be local if it could be mitigated substantially by changes made at the landscape or municipality level. If changes at the watershed or regional level could strongly counteract a driver we classified it as regional to national, and if actions to influence a driver require global or continental coordination we coded it as international. For drivers with management options at more than one scale, we chose the broadest scale at which managerial actions are likely to be strong enough to avoid the shift.

## Network simulations

To better understand the relative importance of regime shifts and drivers we constructed a bipartite network where a driver is connected to a regime shift if there is a reference in the academic literature that suggests causality or influence on its feedback mechanisms. The bipartite network was analysed by considering two network projections: a network of drivers connected by the regime shifts they caused, and a network of regime shifts connected by the drivers they share. Since highly connected drivers are more likely to cause regime shifts and highly connected regime shifts are more vulnerable to different sets of drivers, the mean degree, the co-occurrence index and clustering coefficient^24,25^ were measured and compared with 10000 random simulated networks. We assume that the relative importance of a driver, or the number of times that is reported, depends on our particular sample of regime shifts. Therefore we randomly reshuffled the associations between drivers and regime shifts, keeping the number of links per node unchanged. Simulations were performed in the R statistical software^26^, using a Sequential Importance Sampling algorithm, in R’s networksis^27^ and ergm^28^ packages. The comparison between observed interactions and random data is fundamental to understand whether the co-occurrence patterns are due to sampling noise or corresponds to a real pattern. If the observed patterns deviate from random, there should be theoretical reasons why they diverge that are further explored with statistical modeling.

## Model fitting

Exponential random graph models^29^ were used to explore what local processes could better explain the emergent patterns in the networks. We tested whether certain minimal configurations are statistically more common (e.g. triangles) or if links are significantly more likely to occur if nodes share the same attribute (e.g. management scale). Nestedness^30^ was calculated for the bipartite network to test if the generalist or idiosyncratic character of each driver in the network was related to its scale of management. We used the number of papers reported per regime shift on the ISI Web of Science by 2013 as an approximation of how extensively a regime shift has been studied.

To explore the processes underlying the network patterns, we modelled scale of management, nestedness, frequency and directedness as categorical variables or node covariates for drivers; while ecosystem type, nestedness, number of papers reported, and frequency were modelled as categorical variables or node covariates for regime shifts. The presence or absence of categorical variables in the RSDB was used to construct distance measures of how similar two regime shifts are depending on the variables shared. These distances were modelled as edge covariates for the regime shift network projection (see regime shifts clustering below). The bipartite network was modelled as binary network with geometrically weighted terms^31–33^, while the one-mode projections were modelled following the specifications for weighted edges^34^ and a Poisson distribution as reference. All models were fitted with ergm ^28^ and ergm.count^34^ packages for R^26^.

## Regime shifts and drivers clustering

We used multi-dimensional scaling to investigate the patterns underlying the clustering of regime shifts. First we calculated the Sorensen-Dice distance between regime shifts given the drivers they share. This measure favours the presence of common drivers in the network rather than their absence, and we use it because we are analyzing driver co-occurrence or regime shifts rather than straightforward difference among regime shifts. The hierarchical clustering was performed using the categorical variables of the RSDB after deleting zero columns, grouped by variables as follows: ecosystem processes (5 variables), provisioning services (8), regulating services (8), cultural services (4), drivers (10), land use (11), scales (8), and reversibility (3).

We analysed patterns among the drivers and the regime shifts in two ways, first by using existing classifications from global change research to classify drivers into 5 broad and 15 detailed categories; and second by clustering the drivers based on patterns produced by their connections to regime shifts. Applying matrix multiplication of the bipartite data by the drivers categorization, we obtained the number of drivers per regime shift that fall into each broad and detailed global change category. Euclidean distances were used to organize the drivers into hierarchical clusters with an average method using the R package gplots^35^. These two approaches allowed us to compare how global change meta-drivers impact regime shifts, and to detect emergent patterns from our regime shift data based on the published literature.

## Results

We identified 57 drivers underlying 25 regime shifts (Fig 1). The mean number of drivers per regime shift is 11.2, ranging from a low of 3 for *steppe to tundra* to a high of 22 for *mangrove collapse*. The most frequently reported drivers of regime shifts are *climate change, agriculture* and *fishing*, which are reported as drivers of 19, 17 and 15 regime shifts respectively. There are also 14 idiosyncratic drivers (∼24%) that are unique to specific regime shifts. More than half of the connections between drivers and regime shifts are accounted for by 13 drivers (∼22%). The most frequently co-occuring drivers, understood as the number of regime shifts they jointly drive, are *agriculture, climate change, nutrient inputs, deforestation, greenhouse gases, erosion* and *sea surface temperature*, where each pair occurs together in 10 or more regime shifts. The regime shifts with the greatest number of shared drivers are *marine eutrophication, sea grass collapse, fisheries collapse,* and *kelp transitions*, which have 8 drivers in common.

**Figure 1.**
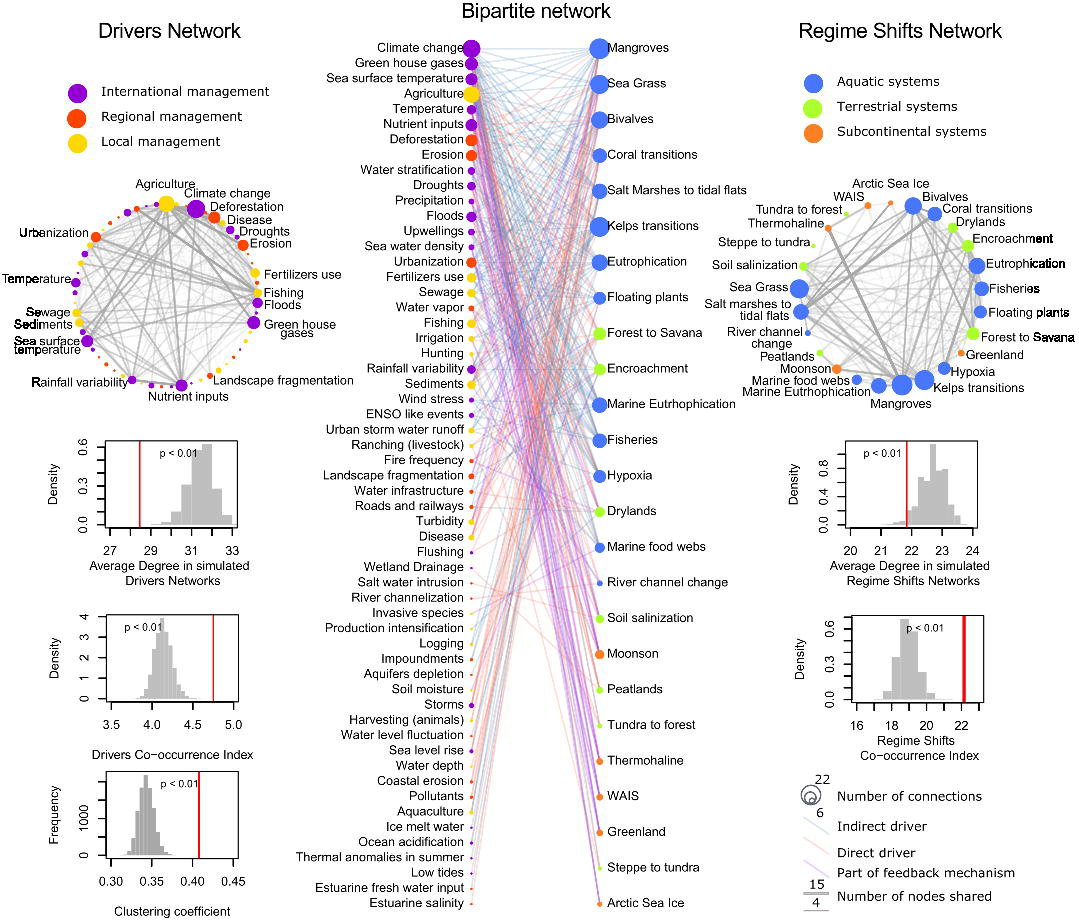
Regime shifts -Drivers Network. In the centre the bipartite network of 57 drivers (left) and 25 regime shifts (right) organized by their nestedness. Highly nested nodes are idiosyncratic and are located on the lower part of the graph while nodes with low nesting are generalist and appear in the upper part. On the right is the one-mode projection of regime shifts (N=25). The width of the links is scaled by the number of drivers shared, while node size corresponds to the number of drivers per regime shift. On the left is the one-mode projection of drivers (N=57), with link width scaled by the number of regime shifts for which causality is shared, and node size proportional to the number of regime shifts per driver. Below each projection is the expected distributions for the co-occurrence index and average degree for the one-mode projection of the drivers and regime shifts networks. The bottom left panel shows the clustering coefficient for the bipartite network. For all structural statistics, the red lines mark the actual values for the observed data.

The regime shift-drivers network had a much higher clustering coefficient, higher co-occurrence index, and lower mean degree than randomized networks (t-test for all statistics P<10^15^, Fig 1). This result suggests that co-occurrence patterns among drivers are related to underlying processes. Furthermore, the network exhibits a nested structure: idiosyncratic drivers co-occur only with drivers that also co-occur with generalist ones (Fig 1 and S1 Fig). Surprisingly, the exponential random graph models show (S2 Table) that the nested structure of the network is not due to global drivers being widely shared among regime shifts and local drivers being idiosyncratic. Rather, drivers that can be managed at local and regional scales are more likely to co-occur with drivers that can also be managed at the same scale. Drivers are significantly more likely to co-occur if they are indirect and generalist. Aquatic and subcontinental regime shifts tend to share the same set of drivers; while terrestrial and subcontinental regime shifts share fewer and more varied sets of drivers. Overall, regime shifts are more likely to share drivers that affect similar ecosystem processes, impact similar ecosystem services, occur in similar ecosystems and occur at similar spatio-temporal scales (S2 Table).

Ecosystem type has a strong influence on the variety of regime shift drivers as well as ecosystem services impacted by regime shifts (Fig 2 & Fig 3). Multi-dimensional scaling reveals that aquatic regime shifts often affect fisheries, water purification, disease control and aesthetic values, and they occur more often at the local scale (S2 Fig). Terrestrial regime shifts are strongly influenced by food production and habitat modification, and surprisingly also by oceanic spillovers. They consistently affect water cycling, the provision of food crops and fresh water, and occur on land uses related to agriculture. Subcontinental regime shifts are quite different in being almost completely driven by anthropogenic greenhouse gases, climate, ecological, and oceanic spillover effects. Interestingly, they consistently affect climate regulation and occur at time scales of centuries. Based upon our classification of regime shift drivers, we found that climate related drivers are shared across all regime shifts, while oceanic and ecological spillovers are shared across the majority of regime shifts. Aquatic regime shifts are driven by all major types of global change drivers, with no drivers related to terrestrial resource extraction or fire (Fig 2). Almost two thirds of the identified regime shift drivers (62%) have the potential to be managed at local or national scales, while a third (38%) can only be managed internationally (Fig 3).

**Figure 2.**
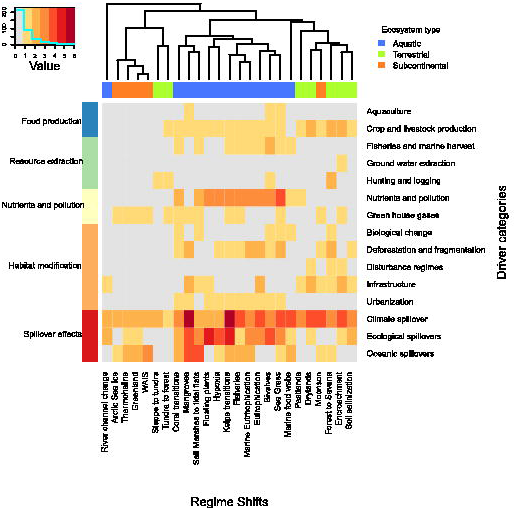
Driver categories per regime shift. Shading intensity indicates the number of drivers per regime shift that falls in each driver category. The dendrogram represents the similarity of regime shifts given the drivers shared (rows) based on hierarchical clustering with an average method upon Euclidean distances. The grey area shows categories with missing drivers. The upper horizontal bar shows the ecosystem type while the left lateral bar shows the 5 broad categories into which the 15 specific drivers categories shown in the rows (right) are classified.

**Figure 3.**
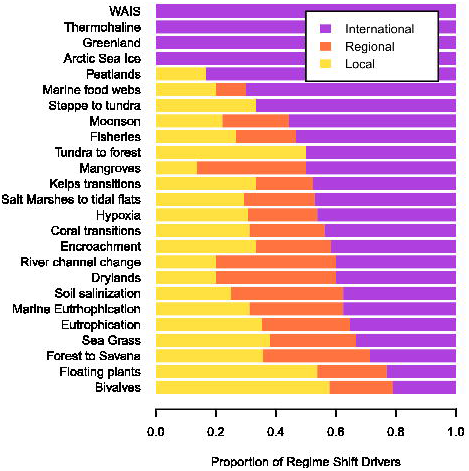
Managerial opportunities per regime shift. Each bar shows the proportion of drivers that can be managed at different scales.

## Discussion

The variety of drivers revealed by our analysis demonstrates that reducing the risk of regime shifts requires integrated action on multiple dimensions of global change across scales (Fig 2 and 3), a non-trivial challenge for governance. Even heroic actions, such as halting climate change or halting agricultural expansion, if not combined with other actions, will be insufficient to avoid most regime shifts.

Food production and climate change are key drivers of regime shifts that are intertwined with one another (Fig 2) and expected to increase in the coming decades^4,36,37^. These drivers have the potential to synchronize the risk of regime shifts across many systems as well as to produce cascading regime shifts. Drivers related to food production consist of a broad set of drivers that tend to occur together. They combine resource extraction (e.g. fishing, cropping), nutrients and pollution and strongly co-occur with habitat modification drivers (e.g. urbanization, deforestation), all of which simplify and homogenize ecosystems. Climate related drivers are a more narrow set of connected drivers, providing few opportunities for local or regional management. However in both cases there is strong potential to reduce risk of synchrony by managing local and national scale drivers^38,39^. Local activities and global markets connect climate and food drivers, which increases the risk of synchronized regime shifts, but also provides an opportunity to increase resilience by diversifying local and national energy, food, and regime shift management.

The number of regime shifts that share climate and food production related drivers furthermore increases the potential for cascading effects among multiple regime shifts. Cascades of regime shifts are possible when some regime shifts enhance the drivers of other types of regime shifts^14,37,40,41^. Regime shifts that contribute to climate change by releasing greenhouse gases or decreasing albedo, or regime shifts that increase the demand for food by e.g. decreasing crop production, can increase the likelihood of other climate or food production driven regime shifts far away.

It remains unclear whether the differences between aquatic, terrestrial and subcontinental regime shifts are explained by the extent to which they have been studied. In the early development of regime shifts theory, aquatic systems were proposed as ideal candidates to test for the existence and mechanisms underlying these non-linear dynamics^15^, and consequently have been better studied. Aquatic environments also have and share more drivers, often accounting for land and ocean interactions. Subcontinental regime shifts are harder to study since most evidence relies on observation of long-term processes rather than experimentation. They also share many drivers but to a lesser extent than aquatic regime shifts, and their drivers and impacts are typically climate related. This makes them ideal candidates for the study of cascading effects, when one regime shift acts as a driver of other shifts. Terrestrial regime shifts tend to have more idiosyncratic drivers. They are also prone to cross-scale interactions, when the aggregation of many instances of the same regime shift scales up to affect drivers that further exacerbate the risk of the regime shift elsewhere. Well studied examples of this effect are percolation thresholds for fire, erosion and landscape fragmentation^40,42,43^.

Reducing local drivers can build resilience to continued global change, but unless the rates of global change are slowed or reversed, these changes will eventually overwhelm local management^44^. Furthermore, our results (S2 table) suggest that in situations where regime shifts and their drivers are poorly understood, managerial options that work for well-understood regime shifts could potentially be applied to uncertain or data scarce regime shifts if they share similar ecosystem processes, impact similar ecosystem services, occur in similar ecosystems and occur at similar spatio-temporal scales. Similarly, our results suggest that while monitoring direct drivers allows change in the risk of a regime shift to be estimated, management efforts are likely more effective when targeting indirect and generalist drivers because these drivers influence many types of regime shifts, and therefore reducing them can reduce the risk of multiple regime shifts.

This paper has presented a novel comparison of regime shifts and their drivers. The development of the regime shift database and the framework for comparison offers a platform for others to extend this work. The regime shifts database framework facilitated comparison of diverse types of regime shifts, broadening our understanding of regime shift similarities at the conceptual level while offering the possibility to translate the observed patterns into useful management insights. Our coding of drivers was done in a systematic, repeatable way, and although some of the categories could have been defined differently, we do not believe it would alter the overall pattern of our results. However, future work needs to take into consideration that the weighting of drivers is not homogeneous across all regime shifts, as such weights are expected to be context dependent. Furthermore, our network approach so far does not allow us to infer the role of dynamics, how changes in the intensity of drivers over time strengthens or weakens their interaction, or how the ordering of events could exacerbate or dampen the effect of such interactions.

Achieving a sustainable future will require meeting needs for ecosystem services^9^, while avoiding regime shifts that disrupt the resilient production of these services. Consequently, both theoretical and empirical work is needed to better assess where regime shifts are most likely to happen, which ecosystems and their services will be most affected, and which groups of society will be most impacted. Furthermore, better understanding of the dynamics of regime shifts and their drivers is needed to understand the i) extent to which increasing drivers of global change can trigger synchronous regime shifts; and ii) how regime shifts, by altering the drivers of other regime shifts, can trigger or inhibit cascades of regime shifts.

## Acknowledgements

We thank contributors of the regime shift database and useful comments on the manuscript by Steve Carpenter, Carl Folke, Line Gordon, Stuart Kininmonth, Will Steffen, Daniel Ospina and Steven Lade.

## Supplementary Information

**S1 File. A regime shift worked example and its causal loop diagram.**

**S1 Figure. Drivers clustering.** Shading intensity indicates the similarity between drivers given the regime shifts they cause. The row dendrogram shows a hierarchical clustering calculated on the Sorencen-Dice distance of the drivers matrix. The column side bar shows the scale of management per driver.

**S2 Figure. Multi-dimensional scaling.** Regime shifts are organized according to their similarity given shared drivers, with a) names coloured according to ecosystem type: blue = marine regime shifts, green = terrestrial and orange = subcontinental regime shifts. Smaller plots show the environmental fitting for subsets of the regime shift categorical variables: b) ecosystem processes (5 variables), c) provisioning services (8), d) regulating services (8), e) cultural services (4), f) drivers (10), g) land use (11), h) scales (8), and i) reversibility (3). Only variables that significantly (p<0.05) influence the regime shifts ordering given their shared drivers are shown in purple as vectors, indicating the directionality of their influence.

**S1 Table.** Summary of the 25 regime shifts examples from the regime shifts database used in this analysis. *Only the main ecosystem service impacts are shown.

| Regime Shift | Initial regime | Alternative regime | Ecosystem | Ecosystem Services affected * |
| --- | --- | --- | --- | --- |
| Eutrophication | Clear water | Murky water | Aquatic - Coastal | - Fisheries<br>- Water purification<br>- Recreation |
| Marine food web simplification | Predators dominated | Lower trophic groups dominated | Aquatic - Coastal | - Fisheries<br>- Pest & disease regulation<br>- Recreation |
| Hypoxia | Normoxia | Hypoxia, anoxia | Aquatic - Coastal | - Fisheries<br>- Pest & disease regulation<br>- Recreation |
| Fisheries collapse | High abundance of commercial fish | Low abundance of commercial fish | Aquatic - Marine | - Fisheries<br>- Pest & disease regulation<br>- Biodiversity |
| Floating plants | Submerged plants dominance | Floating plants dominance | Aquatic | - Fisheries<br>- Pest & disease regulation<br>- Recreation |
| River channel change | Old channel course | New channel regime | Aquatic | - Freshwater<br>- Food production<br>- Regulation soil erosion<br>- Transport |
| Mangroves transitions | Mangrove forest | - Salt marshes<br>- Rocky tidal<br>- Shrimp farms | Aquatic – coastal | - Fisheries<br>- Timber<br>- Regulation soil erosion<br>- Recreation |
| Sea grass transitions | Sea grass dominated | - Algae dominated<br>- Bare sediments | Aquatic – coastal | - Fisheries<br>- Water purification<br>- Regulation soil erosion |
| Marine eutrophication | Clear water | Nutrient rich water | Marine | - Fisheries<br>- Water purification<br>- Recreation |
| West Antarctica Ice Sheet collapse | Full glacial or modern interglacial | Extreme interglacial | Polar | - Climate regulation<br>- Natural hazards protection |
| Bivalves collapse | High abundance of bivalves | Low abundance of bivalves | Marine | - Water purification<br>- Fisheries<br>- Biodiversity |
| Coral transitions | Coral dominated reefs | - Macro-algae<br>- Soft corals<br>- Corallimorpharians<br>- Sponges<br>- Urchin barren | Marine | - Biodiversity<br>- Fisheries<br>- Recreation<br>- Coastal protection |
| Kelp transitions | Canopy forming algae | - Turf forming algae<br>- Urchin barrens | Marine | - Fishing<br>- Biodiversity<br>- Recreation |
| Encroachment | Grass dominated savanna | Shrub dominated savanna | Savannas | - Livestock<br>- Climate regulation<br>- Biodiversity |
| Soil salinization | Low salinity soils | High salinity soils | Dry lands | - Fresh water<br>- Food production<br>- Soil erosion regulation<br>- Biodiversity |
| Forest to savannas | Forest | Savanna | Forest - Savanna | - Biodiversity<br>- Climate regulation<br>- Water cycling<br>- Food production |
| Dry land degradation | Dry lands: savannas, dry forest | Deserts | Dry lands | - Freshwater<br>- Food production<br>- Timber and fuel<br>- Climate regulation<br>- Water regulation |
| Tundra to forest | Tundra | Forest | Tundra | - Livestock<br>- Wildlife food<br>- Climate regulation<br>- Timber |
| Monsoon | Strong monsoon | Weak monsoon | Marine - Terrestrial | - Water cycling<br>- Food production, timber<br>- Climate regulation |
| Peatlands | Low productivity & high C accumulation | High productivity & low C accumulation | Peatlands | - Nutrient cycling (C)<br>- Climate regulation |
| Greenland Ice Sheet melting | Permanent ice sheet | No permanent ice sheet | Polar | - Coastline protection<br>- Climate regulation<br>- Water regulation |
| Thermohaline Circulation Collapse | Strong thermohaline circulation | Collapse of thermohaline circulation | Polar - Marine | - Climate regulation<br>- Biodiversity<br>- Food production |
| Salt marshes to tidal flats | Salt marshes | Tidal or subtidal flat | Marine - coastal | - Pollution filtration<br>- Storm protection<br>- Fisheries, food production. |
| Arctic Sea Ice collapse | Arctic with summer ice | Arctic without summer ice | Polar | - Climate regulation<br>- Aesthetic values<br>- Natural hazards protection |
| Steppe to tundra | Steppe | Tundra | Steppe | - Biodiversity<br>- Food production<br>- Climate regulation |

**S2 Table.**
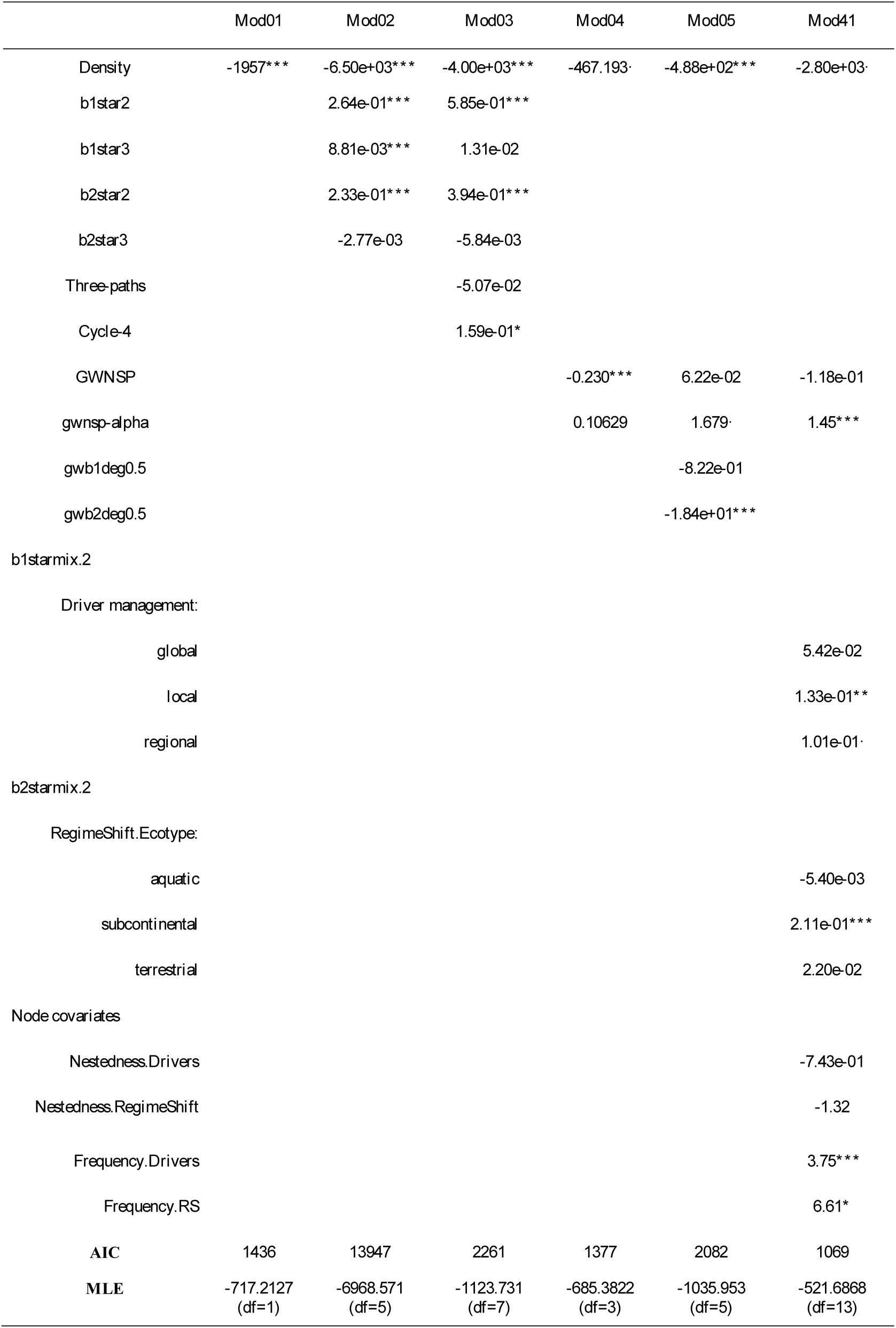
Summary of exponential random graph models fitted to the bipartite and one mode network data.

Models 01 to 05 are null models following the specifications for bipartite networks of ref 42. Model 01 is a Markov random model, model 02 explores the effect of 2 and 3 paths on both projections of the bipartite network, model 03 explore the effects of three-paths and cycles also known as clustering model; model 04 is a curved exponential model that show the effects of geometrically weighted node shared partners (gwnsp), complemented in model 05 by adding geometrically weighted terms for the degree on each one-mode projection. Model 41 is the model that exhibited the best fit following both Akaike Information Criterion (AIC) and Maximum Likelihood Estimation (MLE). Model 41 combines a curved exponential model and explores the effects of homophily – the likelihood of two nodes of being connected on the one-mode projections given that they share attributes: scale of management for driver nodes, ecosystem type of regime shifts nodes, and nestedness and frequency as node covariates respectively. All model are dyadic dependent, only model 41 do not exhibit degeneracy. Significance levels: ***P<0.001, **P<0.01, *P<0.05, .P< 0.1

